# PrimPol variant V102A with altered primase and polymerase activities

**DOI:** 10.1101/2023.11.29.569302

**Authors:** Elizaveta O. Boldinova, Yulia V. Filina, Regina R. Miftakhova, Yana F. Shamsutdinova, A.V. Makarova

## Abstract

PrimPol is a human DNA primase-polymerase which restarts DNA synthesis beyond DNA lesions and non-B DNA structures blocking replication. Disfunction of PrimPol in cells leads to slowing of DNA replication rates in mitochondria and nucleus, accumulation of chromosome aberrations, cell cycle delay, elevated sensitivity to DNA-damaging agents. PrimPol has been suggested to be associated with the development of ophthalmic diseases, elevated mitochondrial toxicity of antiviral drugs and increased cell resistance to chemotherapy. Here, we describe a rare missense PrimPol variant V102A with altered biochemical properties identified in patients suffering from ovarian and cervical cancer. The Val102Ala substitution dramatically reduced both the primase and DNA polymerase activities of PrimPol as well as specifically decreased its ability to incorporate ribonucleotides. We suggest that substitutions in this region would likely distort the active site and affect the catalytic activity of PrimPol.

## Introduction

Human primase/polymerase PrimPol was first identified in 2005 and characterized in 2013 (Iyer et al. 2005; García-Gómez et al. 2013; Wan et al. 2013; Bianchi et al. 2013). PrimPol belongs to the superfamily of archaeo-eukaryotic primases (AEPs) and possesses DNA-primase and DNA-polymerase activities. Unlike human replicative primase (PriS/PriL), which synthesizes RNA-primers, PrimPol preferably synthesizes *de novo* DNA*-*primers using deoxyribonucleotides and starting with initiating ATP ribonucleotide (Martínez-Jiménez et al. 2018). PrimPol is present in both the nucleus and mitochondria and plays an important role in maintaining stability of nuclear and mitochondrial genomes (García-Gómez et al. 2013; Keen, Jozwiakowski, et al. 2014; Mourón et al. 2013). It is assumed that the main function of PrimPol is the restart of DNA synthesis at the sites of blocking DNA damage or secondary DNA structures such as bulky DNA lesions (Piberger et al. 2020), DNA-crosslinks (González-Acosta et al. 2021), G-quadruplexes (Schiavone et al. 2016; T. Li et al. 2023; Butler et al. 2020), R-loops (Šviković et al. 2019). PrimPol is also capable of translesion DNA synthesis and bypasses a number of DNA lesions by direct insertion of nucleotides opposite DNA lesions as well as «skipping» damaged nucleotides (Makarova et al. 2018; E. Boldinova et al. 2022; Stojkovic et al. 2016; García-Gómez et al. 2013; Bianchi et al. 2013; Martínez-Jiménez et al. 2015). Disfunction of PrimPol in the cell leads to slower replication and proliferation, accumulation of chromosomal aberrations and micronuclei, defects in mitochondrial replication, and increased sensitivity of cells to DNA-damaging agents (Wan et al. 2013; Kobayashi et al. 2016; Bailey et al. 2016; Bailey, Bianchi, and Doherty 2019; Mourón et al. 2013).

It was suggested that disfunction of PrimPol contributes to the development of some human diseases, such as hereditary mitochondrial pathologies (e.g., ophthalmic diseases and even muscular dystrophies) (Kasamo et al. 2020; Zhao et al. 2013; Yuan et al. 2020). Comprehensive bioinformatic pan-cancer analysis revealed that aberrant *PRIMPOL* expression or altered protein structure in different types of cancers possibly affect clinical prognosis, cell invasion and anti-tumor immune response (Deng et al. 2023). According to the TCGA PanCancer Atlas Studies (cBioPortal), 72 somatic mutations of *PRIMPOL* gene carrying 59 missense mutations were identified in cancer patients (Cerami et al. 2012). Many mutations have potential to disrupt PrimPol activity since they affect the enzyme active site, binding sites for the PolDIP2 and RPA regulatory proteins, and ZnFn motif (Díaz-Talavera et al. 2022). Several mutations were identified more than in one patient (R76C/H, L96I, N187H/I, K311I/P, P314H/S, E370G/Q, E407K, R417W/Q, Y521S) that makes them promising targets for future studies of association with the cancer risks and disease prognosis (Cerami et al. 2012).

Biochemical study of the properties of the identified PrimPol variants is just began and only few of them are properly characterized up to date. The variants Y89D/N were identified in patients with hereditable high myopia and progressive external ophthalmoplegia as well as healthy individuals (Kasamo et al. 2020; Zhao et al. 2013; Yuan et al. 2020; Keen, Bailey, et al. 2014; J. Li and Zhang 2015). These substitutions reduce affinity to DNA and dNTPs substrates and increase sensitivity of cells to UV radiation (Keen, Bailey, et al. 2014). Several recent studies revealed new inherited autosomal dominant PrimPol mutant variants in patient with high myopia but their biochemical properties are yet to be described: T285A (Haarman et al. 2022), S330C and I369T (Yang et al. 2023). Also a frameshift mutation (V131Gfs*6) resulting from duplication of c.391G of *PRIMPOL* gene was discovered in patients with high myopia in Asian populations (Cai et al. 2019). The somatic variants F522V and I554T found in cancer patients are located in the RPA-binding domain of PrimPol and significantly reduce the ability of PrimPol to bind RPA *in vitro* (Guilliam et al. 2017). The R417W and R417L somatic mutations found in tumors have been shown to greatly impair PrimPol primase activity (Díaz-Talavera 2020). The somatic Y100H substitution detected in lung cancer samples increases rNTP incorporation due to loss of PrimPol ability to discriminate between ribo- and deoxyribonucleotides but slightly decreases dNTP incorporation. The Y100H variant effectively synthesizes DNA *de novo* as the wild-type PrimPol and is able to synthesize RNA primers. It was suggested that enhanced ability of PrimPol Y100H variant to incorporate rNTPs can provide advantages to cancer cells at early stages of carcinogenesis (Díaz-Talavera et al. 2019).

In the present work, we functionally characterized a rare missense PrimPol variant V102A (c.305T>C; rs142122035) found in patients suffering from ovarian and cervical cancer. This variant has been identified in human genome previously (rs142122035; MAF C=0.011223, a total allele frequency from dbGaP aggregated by ALFA) but was not reported as a clinically significant variant. For comparison, we created two mutant PrimPol variants containing substitutions in the active site: D114E and L115M. We provided molecular evidence that the Val102Ala substitution significantly alters the DNA-primase and DNA-polymerase activities of enzyme.

## Materials and Methods

### Protein purification

Mutant variants of human PrimPol were obtained by site-directed mutagenesis. The wild type and mutant GST-tagged PrimPol proteins were purified from Rosetta 2 strain of *E. coli* as described (E. O. Boldinova et al. 2017).

### DNA oligonucleotide substrates

Oligonucleotide substrate containing 8-oxo-G was purchased from The Midland Certified Reagent Company (Midland, USA). Undamaged fluorescent and unlabeled primers and templates were synthesized by Eurogene (Moscow, Russia). Oligonucleotide «p-p-p-12» containing the 5’-terminal adenosine triphosphate was synthesized enzymatically as described (E. O. Boldinova et al. 2023). To prepare the DNA substrate for testing DNA-polymerase activity, Primer-18 was 5’-labeled with [γ-^32^P]-ATP by T4 polynucleotide kinase (SibEnzyme, Russia) and annealed to the corresponding unlabeled Template-55 at a molar ratio of 1:1.1, heated to 75 ºC, and slowly cooled down to 24 ºC. Cy-5 labeled Primer-16 was annealed to the corresponding Template-30 by the same procedure. The sequences of the oligonucleotides used in this study are shown in Table 1.

**Table 1.**
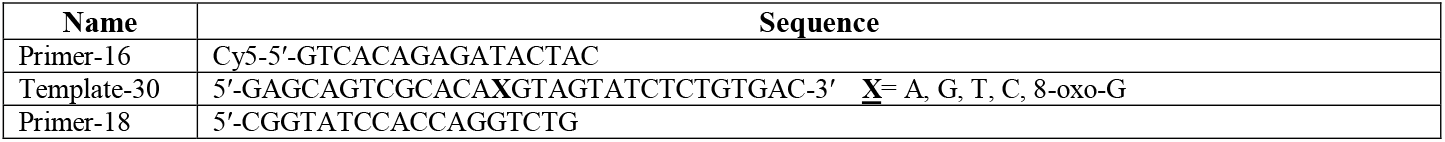
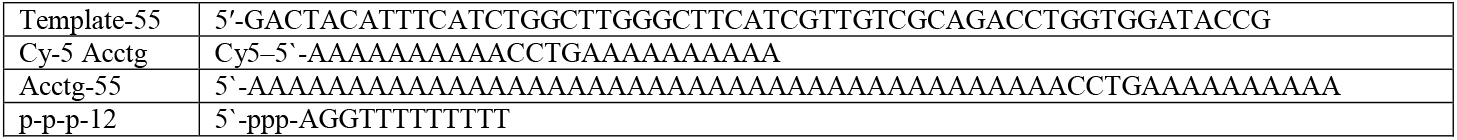
Oligonucleotides used in the study.

### Primer extension reactions

Primer extension reactions were carried out in 20 μl of reaction buffer containing 40 mM HEPES (pH 7.0), 8% glycerol, 0.1 mg/ml BSA, 10 mM MgCl_2_ or 1 mM MnCl_2_, 20-100 nM DNA primer-template substrate, 200 μM dNTPs and 100–800 nM PrimPol. Reactions were started by adding dNTPs and were incubated at 37 °C for specified time. The reactions were terminated by the addition of 20 μl of loading buffer containing 95% formamide, 10 mM EDTA and 0.1% bromophenol blue. DNA products were resolved on 20% PAGE containing 7 M urea, followed by imaging on Typhoon 9400 (GE Healthcare, USA). Experiments were repeated three times. The percent of extended primer (from all DNA bands) was calculated for each reaction. The mean values of primer extension with the standard errors are shown on diagrams.

### Calculation of DNA and RNA polymerization rate constants

To measure the rate of DNA and RNA synthesis, primer extension reactions were stopped after various time intervals (from 2 min to 120 min, as specified in figure legends). Reactions contained 20 nM primer-template DNA, 200 μM PrimPol and 200 μM dNTPs in reactions with Mg^2+^ or 100 nM primer-template DNA, 400 μM PrimPol and 100 μM rNTPs/dNTPs in reactions with Mn^2+^. To calculate the K_obs_ values of the reaction, the kinetics of the primer extension were fit to a single exponential equation: A = A_max_ × [1 − exp(−K_obs_ × t)] using a nonlinear regression, where A is the efficiency of primer extension (the observed % of extended primer), A_max_ is the maximal primer extension, K_obs_ is the observed first-order rate constant for primer extension and t is the reaction time. Calculations were made using GraphPad Prism 8. Experiments were repeated three times.

### Steady-state kinetics analysis of individual dNMP incorporation

To quantify the incorporation of individual dNMPs, we varied each dNTP concentration from 0.3 to 3000 μM in reactions with 100 nM primer-template DNA substrate and 30 nM PrimPol. Reaction mixtures were incubated for different time intervals (1 – 10 min for the wild-type PrimPol, 10 – 60 min for the V102A PrimPol variant, 2 – 20 min for the L115M PrimPol mutant variant). The data were fit to the Michaelis-Menten equation V = V_MAX_ × [dNTP])/(K_M_ + [dNTP]) using a nonlinear regression in GraphPad Prism 8 software, where V and V_MAX_ is the observed and the maximum rates of the reaction (in % of utilized primer per 1 min), respectively, and K_M_ is the apparent Michaelis constant. Calculated apparent K_M_ and V_MAX_ parameters were used to determine the catalytic efficiency (V_MAX_/K_M_) and the fidelity of dNTP incorporation F_inc_ (V_MAX_/K_M_ for non-complementary dNTP divided by V_MAX_/K_M_ for the complementary substrate). Experiments were repeated three times.

### *De novo* DNA synthesis

DNA primase activity was tested in 6 μl reaction mixtures containing 40 mM HEPES pH 7.0, 8% glycerol, 50 μg/ul BSA, 1 mM MnCl_2_, 2 μM unlabeled oligonucleotide Acctg-55 DNA substrate, 200 μM each of dGTP, dCTP and dTTP or dGTP alone, 10 μM dATP, 30 nM [γ-^32^P]-ATP and 2 μM PrimPol. Test tubes were preincubated on ice for 5 min and reactions were started by dNTPs and incubated at 30 ºC for 60 min or for the indicated time intervals. The reactions were stopped by adding an equal volume of loading buffer. Experiments were repeated 2–4 times. The synthesized DNA products were separated on denaturing 30% PAGE with 7 M urea in 1xTBE and visualized on a Typhoon 9400 (GE Healthcare, USA).

### EMSA

Binding of PrimPol to the DNA template-primer substrate with p-p-p-12 annealed to Cy5-Acctg24 was performed in a 20 μl reaction mixture containing 40 mM HEPES pH 7.0, 50 mM KCl, 1 mM DTT, 5% glycerol, 0.1 mg / ml BSA, 1 mM MnCl_2_, 300 nM DNA substrate, and 300–3000 nM PrimPol. Some reactions were supplemented with 1 mM ATP and 200 μM of each dNTP as indicated in the figure legends. The reactions were incubated at 24 ºC for 20 min and placed on ice. Reaction mixtures were directly applied to a 5% native PAAG, and complexes were separated from free DNA in 0.5x Tris-glycine buffer (12.5 mM Tris, 96 mM glycine, pH 8.3) at 10 V / cm and 4 ºC. The gel was visualized on a Typhoon 9400 (GE Healthcare, USA).

### DNA isolation and sequencing

Detection of *PRIMPOL* variants was carried out by targeted sequencing of genomic DNA from 207 patients with malignant epithelial tumors (24-80 years) and 36 healthy volunteers (23-76 years) without a family history of cancer. Samples were obtained from ovarian (n=107), breast (n=79) and cervical (n=21) cancer patients. DNA of 86 ovarian cancer patients was isolated from paraffin-embedded normal tissue with Quick-DNA FFPE Miniprep kit (Zymo Research, USA); other DNA samples were isolated from blood using PureLink Genomic DNA Mini Kit (ThermoFisher Scientific, USA). Specimens were obtained under the local ethical committee-approved protocol (2020-02-13, #8) of the Kazan Federal University. Custom targeted DNA NGS panel was designed covering coding regions of 21 DNA-repair and replication genes including *PRIMPOL* in Ion AmpliSeq Designer (v. 7.02, ThermoFisher Scientific, USA). Amplicon size was 175 bp, coverage 94.89%, target size 86.74 kb. GenSeq reagents (ANPRO, Russia) were used for library preparation. Samples were sequenced using NextSeq 500 with Mid output 300 cycles kit (Illumina, USA). Variations were confirmed by capillary sequencing using self-designed primers (Supplementary Table S1) and BigDye terminator kit v. 3.1. on 3730 DNA Analyzer (ThermoFisher Scientific, USA).

## Results

### Identification of PrimPol V102A variant in patients with cancer

Targeted sequencing of repair and replication genes in 246 cancer patients and healthy volunteers identified PrimPol variant with substitution Val102Ala in the active site of the enzyme. The PrimPol V102A variant was identified in three patients in heterozygosity: one patient in each group with ovarian, breast and cervical cancer (a missense variant in patient with breast cancer was not verified by Sanger sequencing due to lack of DNA material). The aliphatic Val102 residue is located next to the Tyr100 residue, which plays a key role in ribonucleotide discrimination during catalysis (Díaz-Talavera et al. 2019). Since the Y100H (Díaz-Talavera et al. 2019) substitution affects the catalytic activity of PrimPol, mutations of nearby residues can potentially affect properties of PrimPol. For comparison, we also obtained the D114E substitution which affects the key catalytic residue Asp114 coordinating metal ions during catalysis, and the L115M substitution of bulky aliphatic Leu residue located next to the catalytic residues Asp114 and Glu116.

### PrimPol variants with substitutions in the active site have decreased catalytic activity

PrimPol variants were obtained by site-directed mutagenesis and purified from *E. coli* (Figure S1). The DNA polymerase and TLS activities, DNA primase activity and DNA-binding ability of PrimPol variants were analyzed. The D114E substitution resulted in the loss of the DNA primase and DNA polymerase activities regardless the metal ion cofactor used in reactions (Figures 1, 2 and S2). The V102A and L115M substitutions significantly decreased the DNA polymerase activity in the presence of Mg^2+^ ions as a reaction cofactor (Figure 1A, lanes 8-13 and 20-25, 1B).

**Figure 1.**
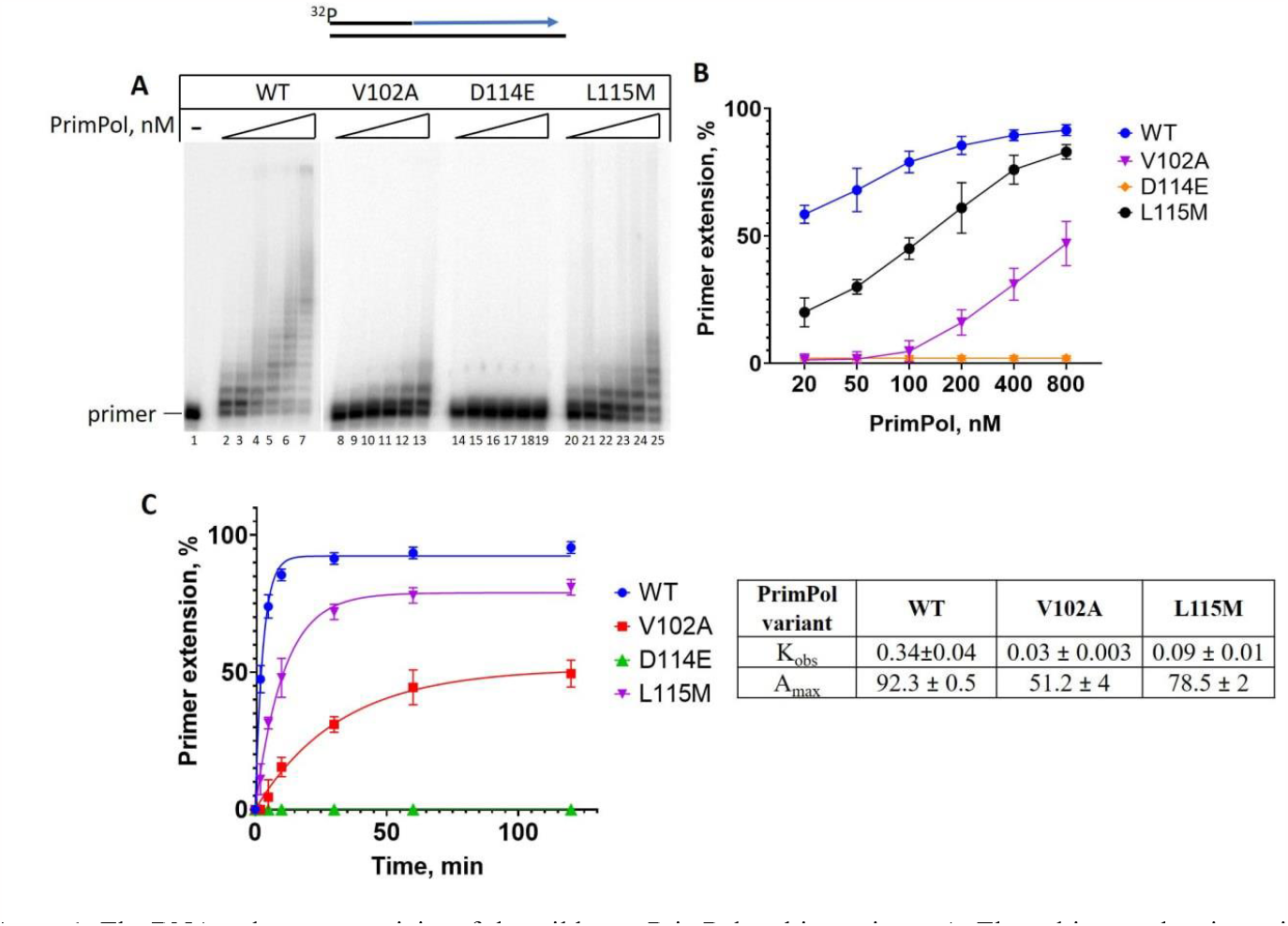
The DNA-polymerase activity of the wild-type PrimPol and its variants. **A**. The gel image showing primer extension by the wild-type PrimPol and its variants on undamaged primer-template DNA-substrate. Reactions were carried out in the presence of 10 mM MgCl_2_, 200 μM dNTPs, 20/50/100/200/400/800 nM PrimPol, 20 nM ^32^P-labeled DNA-substrate for 10 min. **B**. The graph displaying a dependence of DNA polymerase activity on PrimPol concentration in the reactions (shown on panel A). **C**. The graph displaying dependence of DNA synthesis on reaction time and K_obs_ values. Reactions were carried out in the presence of 10 mM MgCl_2_, 200 μM dNTPs, 200 nM PrimPol, 20 nM ^32^P-labeled DNA-substrate for 2/5/10/30/60/120 min. Experiments were repeated three times, the mean values and standard errors are presented.

**Figure 2.**
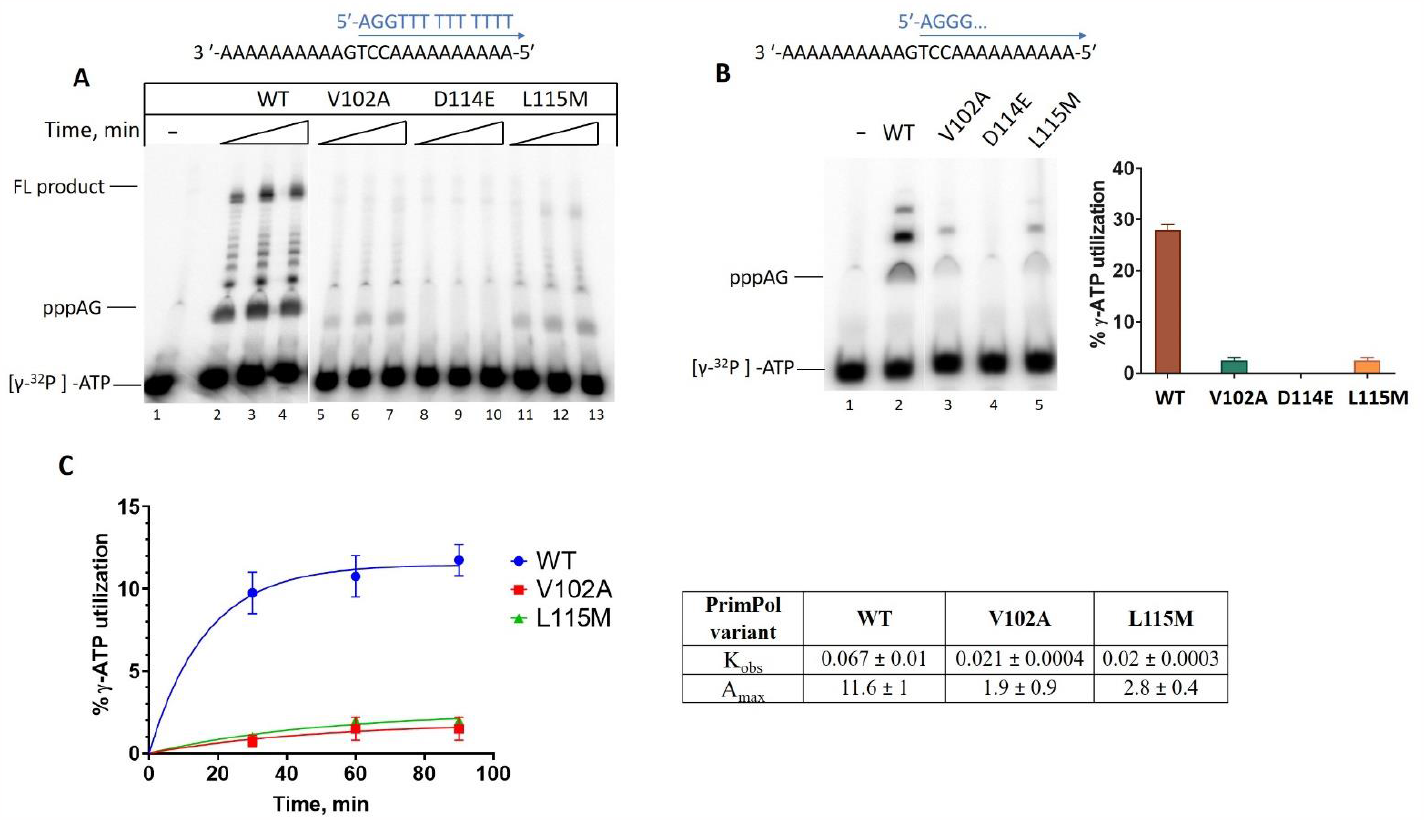
Effect of mutations on the DNA-primase activity of PrimPol. **A**. Comparison of the DNA-primase activity of PrimPol and its variants. Reactions were incubated in the presence of [γ-^32^P]-ATP, ATP, dGTP and dTTP, 2μM PrimPol and 2 μM ssDNA, pH 7.0 and 1 mM Mn^2+^ for 30/60/90 min. **B**. The dinucleotide formation by PrimPol variants. Reactions were incubated in the presence of [γ-^32^P]-ATP, ATP and dGTP, 2 μM PrimPol and 2 μM ssDNA, pH 7.0 and 1 mM Mn^2+^ for 30 min. The obtained results are shown as a bar graph below. **C**. The graph displaying a dependence of DNA primase activity on reaction time and K_obs_ values. Reactions were incubated in the presence of [γ^-32^P]-ATP, ATP, dGTP and dTTP, 2 μM PrimPol and 2 μM ssDNA, pH 7.0 and 1 mM Mn^2+^ for 30/60/90 min. Experiments were repeated three times, the mean values and standard errors are presented.

PrimPol variants synthesized much shorter DNA products than the wild-type PrimPol even at high concentration of enzyme. The time course of primer extension reactions also showed that the substitutions significantly decreased the DNA polymerase activity of PrimPol (Figure 1C) reducing the catalytic rates of PrimPol (in reactions with Mg^2+^) about 11-fold and 3.7-fold for the V102A and L115M PrimPol variants, respectively (Figure 1C). The DNA polymerase activity of mutant proteins was higher in the presence of Mn^2+^ ions but still did not reach the level of the wild-type protein activity (Figure S2).

The DNA primase activity of PrimPol is greatly stimulated by Mn^2+^ ions. Investigation of DNA *de novo* synthesis was carried out in reactions containing ssDNA-template with the «GTCC» primase-binding site, [γ^-32^P]-ATP as initiating nucleotide and rATP/dGTP/dTTP or rATP/dGTP for analysis of total primase activity or dinucleotide formation, respectively. The DNA primase activity of the V102A and L115M PrimPol variants significantly decreased (Figure 2A, lanes 5-7 and 11-13) due to the inhibition of dinucleotide formation (Figure 2B, lanes 3 and 5, 2C). The catalytic rates of PrimPol V102A variant decreased by approximately 3-fold for the DNA primase activity (Figure 2C) and 2.6-fold for the DNA polymerase activity (Figure 1C), in the presence of Mn^2+^ ions. The catalytic rate of the L115M PrimPol variant decreased also by 3-fold for the DNA primase activity (Figure 2C).

The DNA-binding ability of all variants was tested by EMSA with a dsDNA substrate containing a primer with adenosine triphosphate at the 5’-end (E. O. Boldinova et al. 2023). The effect of the V102A and D114E substitutions on DNA binding ability of PrimPol was not statistically significant but the L115M substitution slightly decreased DNA binding of PrimPol although Leu115 residue is located next to the catalytic residues but not the DNA-binding residues (Figure S3). A small effect of these substitutions on affinity to DNA can be explained by a fact that the C-terminal domain plays a main role in template:primer binding (E. O. Boldinova et al. 2023).

To analyze the fidelity of DNA synthesis by the V102A and L115M PrimPol variants we performed single nucleotide incorporation experiments in the presence of Mg^2+^ or Mn^2+^ ions under conditions ensuring approximately equal DNA polymerase activity level between the wild-type enzyme and PrimPol variants. For this reason, we adjusted the protein concentration and the reaction times as indicated in the figure legends. The V102A and L115M substitutions did not show significant effect on the DNA synthesis fidelity in the presence of neither Mg^2+^ (Figure S4) nor Mn^2+^ ions (Figure 3B). The V102A and L115M substitutions slightly decreased incorporation of complementary nucleotides: V102A – opposite A and T (p<0.05); L115M – opposite A and G (insignificantly), T and C (p<0.05). Small increase of dCMP incorporation opposite non-complementary bases in the presence of Mn^2+^ ions was observed only for the L115M PrimPol variant (Figure 3A, 3B).

**Figure 3.**
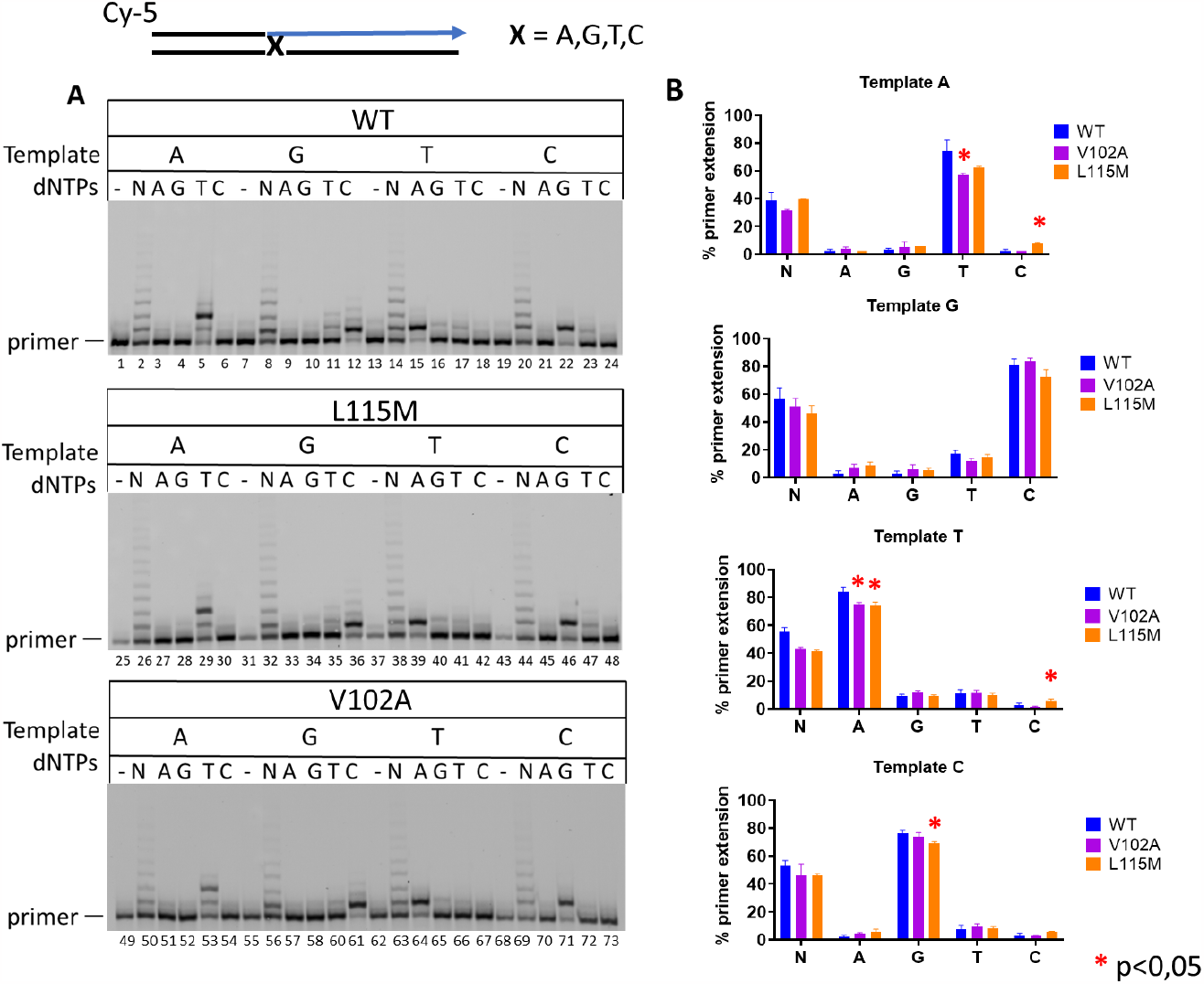
Individual dNMPs incorporation by PrimPol variants in the presence of Mn^2+^ ions. **A**. The gel image showing primer extension by PrimPol and its V102A and L115M variants on undamaged primer-template DNA-substrates. **B**. The diagram showing the percentages of single dNMP incorporation. The reactions were carried out in the presence of 1 mM MnCl_2_, 100 nM Cy5-DNA substrate, 200 μM of dNTP or individual dATP, dGTP, dTTP, dCTP and 200 nM wild-type PrimPol (2 min), 400 nM V102A variant (5 min), 200 nM L115M variant (7 min). The experiments were repeated 2 – 4 times, the mean values and standard errors are presented.

The TLS activity of the V102A and L115M PrimPol variants was tested on the primer-template DNA-substrate containing 8-oxo-G lesion under conditions ensuring equal activity levels of enzymes demonstrated in the reactions on undamaged control DNA. In the single nucleotide incorporation assay in the presence of Mg^2+^ ions, we observed the decreased misincorporation of dAMP opposite 8-oxo-G by both variants while incorporation of complementary dCMP and overall effectiveness of TLS opposite 8-oxoG did not change significantly (Figure 4A, 4C). In the reactions with Mn^2+^ ions, we observed a decrease of dAMP misincorporation for the V102A PrimPol variant, but not for the L115M substitution (Figure 4C).

**Figure 4.**
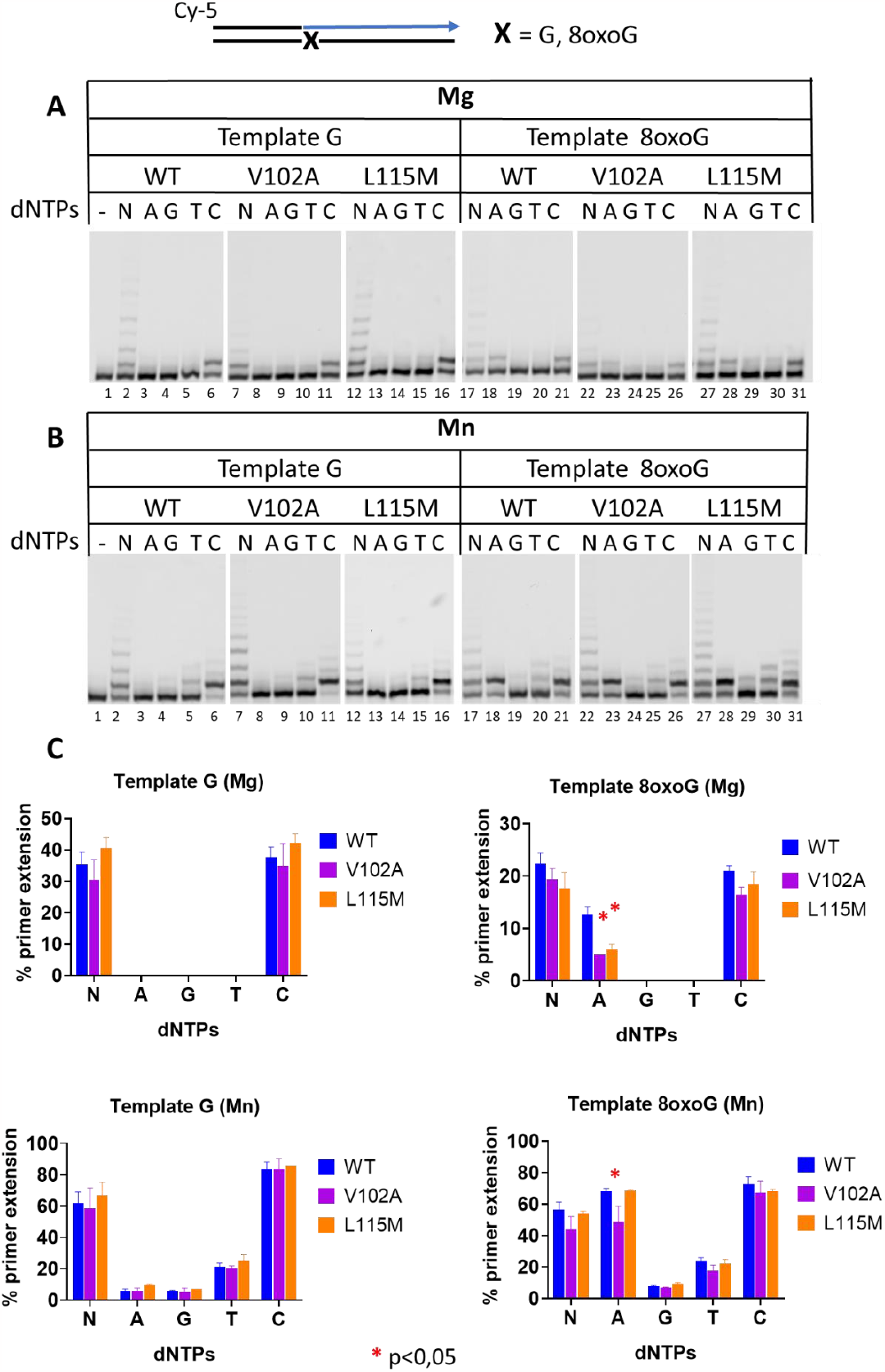
The TLS activity of PrimPol variants opposite 8-oxo-G lesion. **A**. The gel image showing primer extension by the wild-type PrimPol and its V102 and L115M variants on primer-template DNA-substrate containing 8-oxo-G in the presence of Mg^2+^ ions. The reactions were carried out in the presence of 10 mM MgCl_2_, 100 nM Cy5-DNA substrate, 200 μM of dNTP or individual dATP, dGTP, dTTP, dCTP and 200 nM wild-type PrimPol (4 min), 600 nM V102A variant (90 min), 200 nM L115M variant (30 min). **B**. The gel image showing primer extension by the wild-type PrimPol and its V102 and L115M variants on DNA with 8-oxo-G in the presence of Mn^2+^ ions. The reactions were carried out in the presence of 1 mM MnCl_2_, 100 nM Cy5-DNA substrate, 200 μM of dNTP or individual dATP, dGTP, dTTP, dCTP and 200 nM wild-type PrimPol (2 min), 600 nM V102A variant (5 min), 200 nM L115M variant (7 min). **C**. The diagrams showing the efficiency of single dNMP incorporation by the wild-type PrimPol and its V102 and L115M variants opposite G and 8-oxo-G lesion. The experiments were repeated 2 – 4 times, the mean values and standard errors are presented.

More accurate analysis of kinetic parameters of the reactions revealed that substitutions V102A and L115M decreased efficiency of complementary dCMP incorporation opposite 8-oxo-G lesion to the same extent as opposite template G in the presence of Mg^2+^ (Table 2) and Mn^2+^ ions (Table S2). Effect of amino acid substitutions on dAMP incorporation opposite 8-oxo-G was more prominent. In particular, in reactions with Mg^2+^, complementary dCMP was incorporated 2-fold and 3-fold more efficient than dAMP by the wild type and by mutant variants, correspondingly (Table 2). Thus, the V102A and L115M substitutions slightly decreased misincorporation of non-complementary dAMP by did not affect significantly incorporation of complementary dCMP opposite 8-oxo-G lesion. In the presence of Mn^2+^ ions, the L115M PrimPol variant demonstrated the same ratio of dCMP and dAMP incorporation opposite 8-oxo-G as the wild-type enzyme, but the V102A substitution slightly decreased dAMP misincorporation opposite the lesion relative to dCMP, thus increasing fidelity (Table S2).

**Table 2.**
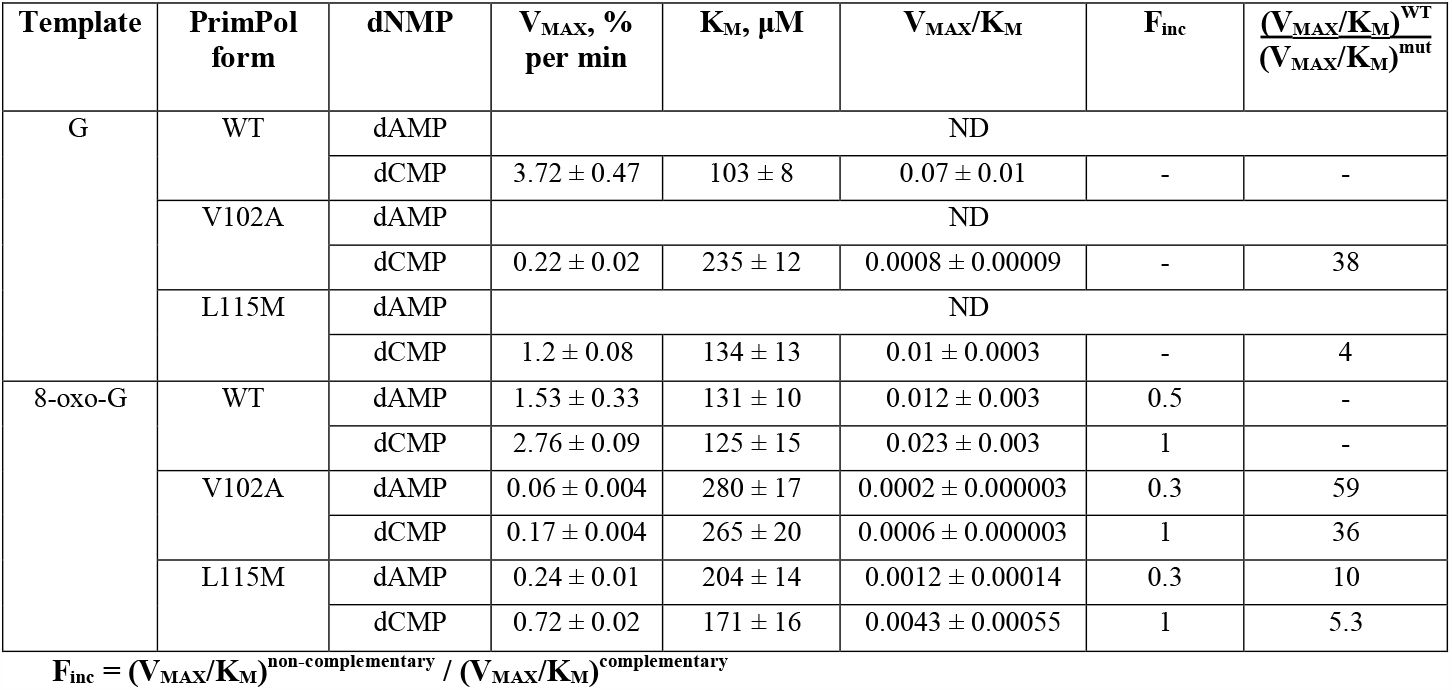
Steady-state kinetic parameters for dNMP incorporation on DNA substrates with G and 8-oxo-G in the +1 position by the wild-type PrimPol, V102A and L115M variants in the presence of Mg^2+^ ions.

| Template | PrimPol form | dNMP | $V_{MAX}$, % per min | $K_M$, $\mu M$ | $V_{MAX}/K_M$ | $F_{inc}$ | $\frac{(V_{MAX}/K_M)^{WT}}{(V_{MAX}/K_M)^{mut}}$ |
| --- | --- | --- | --- | --- | --- | --- | --- |
| G | WT | dAMP | ND |  |  |  |  |
| | | dCMP | $3.72 \pm 0.47$ | $103 \pm 8$ | $0.07 \pm 0.01$ | - | - |
|  | V102A | dAMP | ND |  |  |  |  |
| | | dCMP | $0.22 \pm 0.02$ | $235 \pm 12$ | $0.0008 \pm 0.00009$ | - | 38 |
|  | L115M | dAMP | ND |  |  |  |  |
| | | dCMP | $1.2 \pm 0.08$ | $134 \pm 13$ | $0.01 \pm 0.0003$ | - | 4 |
| 8-oxo-G | WT | dAMP | $1.53 \pm 0.33$ | $131 \pm 10$ | $0.012 \pm 0.003$ | 0.5 | - |
| | | dCMP | $2.76 \pm 0.09$ | $125 \pm 15$ | $0.023 \pm 0.003$ | 1 | - |
| | V102A | dAMP | $0.06 \pm 0.004$ | $280 \pm 17$ | $0.0002 \pm 0.000003$ | 0.3 | 59 |
| | | dCMP | $0.17 \pm 0.004$ | $265 \pm 20$ | $0.0006 \pm 0.000003$ | 1 | 36 |
| | L115M | dAMP | $0.24 \pm 0.01$ | $204 \pm 14$ | $0.0012 \pm 0.00014$ | 0.3 | 10 |
| | | dCMP | $0.72 \pm 0.02$ | $171 \pm 16$ | $0.0043 \pm 0.00055$ | 1 | 5.3 |

### PrimPol variant V102A is unable to insert ribonucleotides

Since the V102A substitution is located next to the steric gate Tyr100 residue, we analyzed efficiency of rNMP incorporation by the V102A PrimPol variant. Primer extension reactions were carried out with increasing concentrations of rNTP or dNTP as control. In the presence of Mg^2+^ ions, PrimPol did not incorporate rNMP regardless of nucleotide concentration (Figure S5). Therefore, comparison was conducted in the presence of Mn^2+^ ions under conditions supporting approximately equal DNA polymerase activity of the wild-type enzyme and V102A PrimPol variant. It was shown that the V102A PrimPol variant almost completely lost ability to insert rNMPs (Figure 5). In contrast, the L115M PrimPol variant incorporated rNTP with lower efficiency than the wild-type PrimPol but decrease of activity was not dramatic (Figure 5).

**Figure 5.**
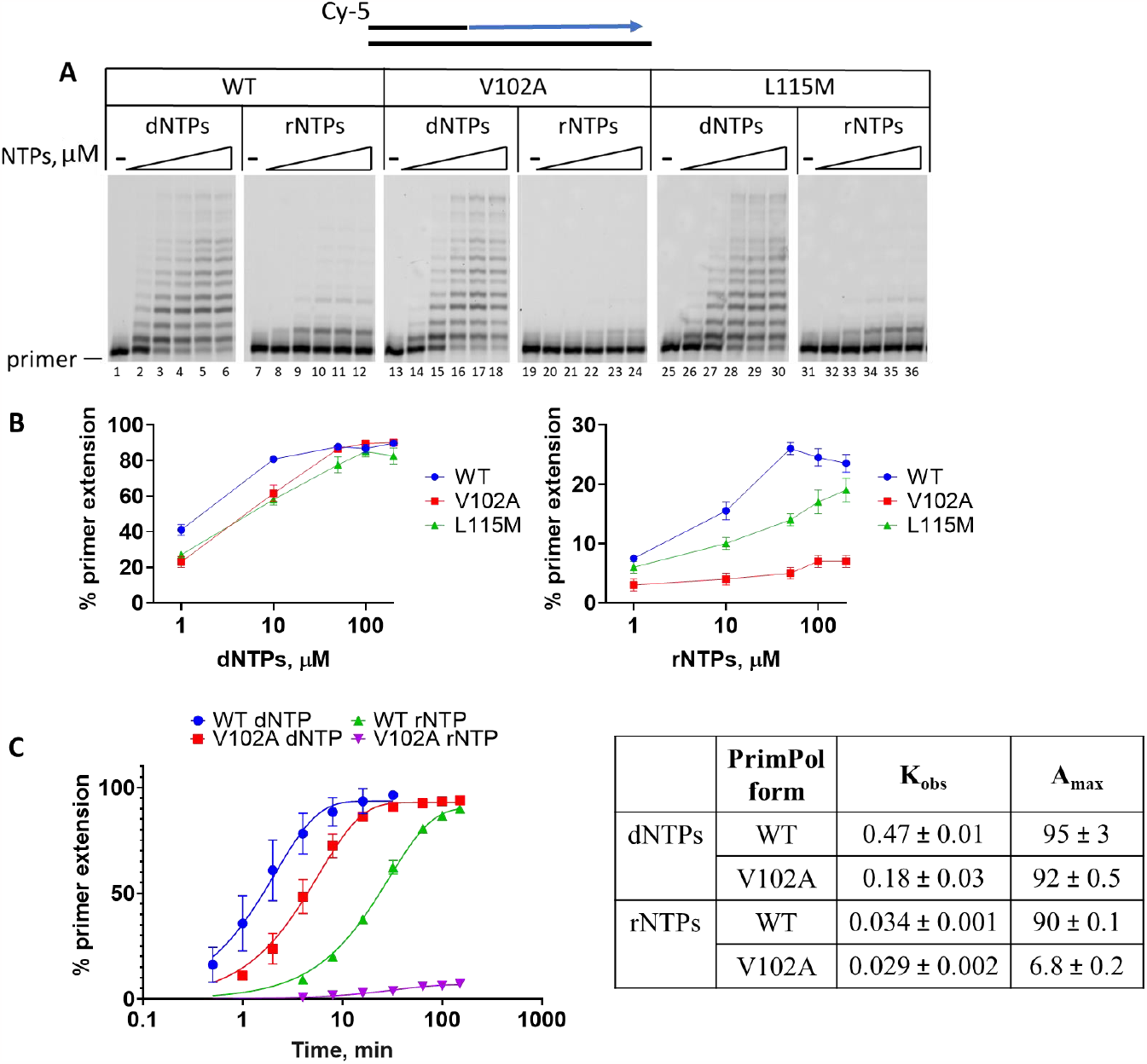
Ribonucleotides incorporation by PrimPol and its V102A and L115M variants. **A**. The gel image showing primer extension by the wild type PrimPol and its V102 and L115M variants using dNTP and rNTP in the presence of Mn^2+^ ions. Reactions were carried out with 100 nM primer-template Cy5-DNA substrate, 1/10/50/100/200 μM dNTPs or rNTPs, 200 nM wild-type PrimPol (10 min), 400 nM V102A variant (20 min), 300 nM L115M variant (20 min), 1 mM MnCl_2_, pH 7.0. **B**. The diagrams showing efficiency of dNTP and rNTP incorporation. **C**. The graph displaying dependence of DNA synthesis efficiency on reaction time and K_obs_ values. Reactions were incubated in the presence of 100 nM Cy5-DNA substrate, 400 nM wild-type PrimPol or V102A variant, 100 μM rNTPs, pH 7.0 and 1 mM Mn^2+^ for 0.5/1/2/4/8/16/32/64/100/150 min. Experiments were repeated two times, the mean values and standard errors are presented.

## Discussion

Targeted sequencing of repair and replication genes revealed a PrimPol variant V102A with substitution in the active site of the enzyme only in patients with cancer. However, our group size was too small to determine the statistical significance of this finding. According to the dbSNP report, it is a rare missense SNP variant with clinical significance status “Not Reported in ClinVar”. Heterozygous substitution V102A is represented in European population with MAF C=0.012371, is very rare in African and Latin American populations (MAF C=0.001–0.002) and absent in Asian population.

Substitutions of aliphatic residues located near key catalytic residues can significantly affect enzymatic activity. The aliphatic Val102 residue is located next to the Tyr100 residue, which forms contacts with the incoming nucleotide substrate and is responsible for the discrimination of deoxy- and ribonucleotides in the active site of PrimPol (Díaz-Talavera et al. 2019). Reducing the size of the side chain by replacing Val102 by Ala may distort the active site affecting the geometry and coordination of bonds formed with the incoming nucleotide, resulting in a decrease in the efficiency of nucleotide incorporation. Indeed, we demonstrated that the V102A substitution did not affect binding of PrimPol to DNA but significantly decreased its DNA primase and DNA polymerase activities as well as dramatically reduced ability to incorporate ribonucleotides. This reduction affected the DNA primase and DNA polymerase activities to the same extent (2.6–3-fold in reactions with Mn^2+^) suggesting that the V102A substitution altered catalysis during the elongation step. It is likely that the V102A substitution increases PrimPol selectivity toward dNTPs due to strengthening of the steric gate for rNTPs mediated by Tyr100. In the V102A mutant, Tyr100 can move closer to the C2 of a sugar of incoming NTP and make a stronger clash with 2’OH of a ribose.

It is important to note that mutations nearby of Tyr100 were suggested to be possibly involved in the progression of lung cancer (Díaz-Talavera et al. 2019; Liu et al. 2012). In particular, the Y100H substitution can facilitate ribonucleotide incorporation promoting cell proliferation under conditions of dNTP depletion at the early stages of carcinogenesis (Díaz-Talavera et al. 2019). We discovered that the V102A variant demonstrated the opposite phenotype inhibiting rNMP incorporation, and the substitution of nearby residue Leu115 also decreased rNMP incorporation. Thus, the entire 100-115 a.a. region of PrimPol is important for nucleotide coordination and mutations and polymorphic substitutions in this region may affect PrimPol functioning.

Deficiency in PrimPol might be associated with ophthalmological diseases (Kasamo et al. 2020; Yuan et al. 2020; Haarman et al. 2022; Yang et al. 2023; Cai et al. 2019), altered response to DNA damage and chemotherapy (Quinet et al. 2019), increased toxicity of retroviral nucleotide analog therapy (Duong et al. 2020) as well as increased cancer suspensibility (Deng et al. 2023). Deletions of *PRIMPOL* were found to be common for breast cancer (Ciriello et al. 2015). Altogether, our data demonstrated that the PrimPol V102A polymorphic variant possesses dramatically altered catalytically properties likely affecting PrimPol functions *in vivo*. The clinical effect of V102A variant requires future investigation.

The D114A substitution, affecting a key residue coordinating metal ions in the active site, abrogates catalytic activity of PrimPol (Keen, Jozwiakowski, et al. 2014; Rechkoblit et al. 2016; Calvo et al. 2019). We demonstrated that the Asp114 substitution to the functionally similar Glu residue also led to complete loss of catalytic activity in the presence of both Mg^2+^ and Mn^2+^ ions. Replacing the aspartate with glutamate does not change the overall charge, but should affect a coordination of catalytic metals and dNTP binding. It was previously shown that a reverse substitution of the second key catalytic residue coordinating metal ions, E116D, disrupts the binding of metal ions by PrimPol and suppresses the activity of the enzyme (Calvo et al. 2019).

Moreover, we also demonstrated that substitution of bulky aliphatic Leu115 adjacent to the key catalytic residues Asp114 and Glu116 in the active site of PrimPol also leads to decrease in the DNA-primase and DNA-polymerase activities. It can be assumed that the replacement of leucine with a bulkier methionine leads to local destabilization of protein fold and disturbs the active site. These data highlight the importance of a stable active site conformation for efficient catalysis and predict that substitutions in this region would likely affect PrimPol activity. Altogether, these data provide the first functional molecular characterization of a missense PrimPol V102A variant and shed light on the mechanism of DNA synthesis in the active site of PrimPol as well as predict critical residues that may have clinical significance.

## Acknowledgments

We thank Andrey Baranovskiy (UNMC, Omaha, USA) for helpful discussion and suggestions.

