## Supplemental material for "PrimPol variant V102A with altered primase and polymerase activities"

*[3] "Translational oncology" research laboratory, Institute of Fundamental Medicine and Biology, Kazan Federal University, Kremlevskaya 18, 420008 Kazan, Russia*

*[4]Chemotherapy department №1, Republican Clinical Oncology Dispensary of the Ministry of Health of the Republic of Tatarstan named after prof. M.Z.Sigal, Sibirskiy trakt 29, 420029 Kazan, Russia*

**Table S1**. Primers for Sanger sequencing

| Target | Primer sequence 5'→3' | |
|---|---|---|
| c.305T>C (p.V102A)<br>c.342T>G (p. D114E)<br>c.343T>A (p. L115M) | Forward | TCAGTCTCAGTCAGTCAATCAATC |
| | Reverse | AGCTATTCGTAATGGAAGTCAGAG |

**Table S2**. Steady-state kinetic parameters for dNMP incorporation on DNA substrates with G and 8-oxo-G in the +1 position by the wild-type PrimPol, V102A and L115M variants in the presence of $Mn^{2+}$ ions

| Template | PrimPol form | dNMP | $V_{MAX}$, % per min | $K_M$, µM | $V_{MAX}/K_M$ | $F_{inc}$ | $\frac{(V_{MAX}/K_M)^{WT}}{(V_{MAX}/K_M)^{mut}}$ |
|---|---|---|---|---|---|---|---|
| **Mn** | | | | | | | |
| G | WT | dAMP | 1.1 ± 0.04 | 10.2 ± 1.1 | 0.11 ± 0.02 | 0.02 | - |
| | | dCMP | 19.8 ± 0.1 | 4.2 ± 1 | 4.8 ± 1.2 | 1 | - |
| | V102A | dAMP | 0.1 ± 0.01 | 44.2 ± 1.9 | 0.003 ± 0.00005 | 0.003 | 42 |
| | | dCMP | 10.34 ± 1.6 | 14.3 ± 5 | 0.7 ± 0.1 | 1 | 6.4 |
| | L115M | dAMP | 0.3 ± 0.06 | 15.6 ± 5.7 | 0.019 ± 0.003 | 0.02 | 5.6 |
| | | dCMP | 4.9 ± 0.6 | 4.4 ± 0.8 | 1.1 ± 0.4 | 1 | 4 |
| 8-oxo-G | WT | dAMP | 13.61 ± 1.2 | 10.05 ± 2.4 | 1.4 ± 0.33 | 0.5 | - |
| | | dCMP | 15 ± 1.4 | 5.2 ± 1.6 | 3.1 ± 0.8 | 1 | - |
| | V102A | dAMP | 2.2 ± 0.2 | 13.25 ± 1.8 | 0.17 ± 0.04 | 0.4 | 8.5 |
| | | dCMP | 3.5 ± 0.9 | 9.2 ± 1.9 | 0.4 ± 0.02 | 1 | 8 |
| | L115M | dAMP | 3.35 ± 0.4 | 9.4 ± 1.3 | 0.36 ± 0.09 | 0.5 | 3.8 |
| | | dCMP | 4.8 ± 0.8 | 5.6 ± 0.5 | 0.7 ± 0.01 | 1 | 4.6 |

$F_{inc} = (V_{MAX}/K_M)^{non\text{-}complementary} / (V_{MAX}/K_M)^{complementary}$

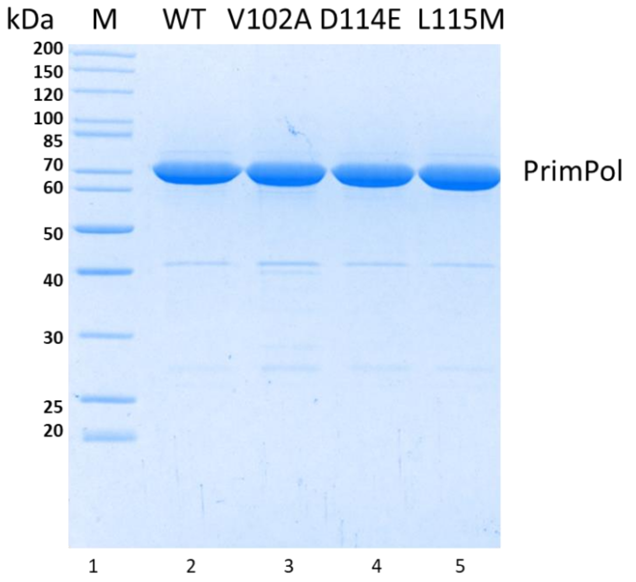


**Figure S1**. The SDS-PAGE of purified wild-type PrimPol and itsV102A, D114E and L115M variants.

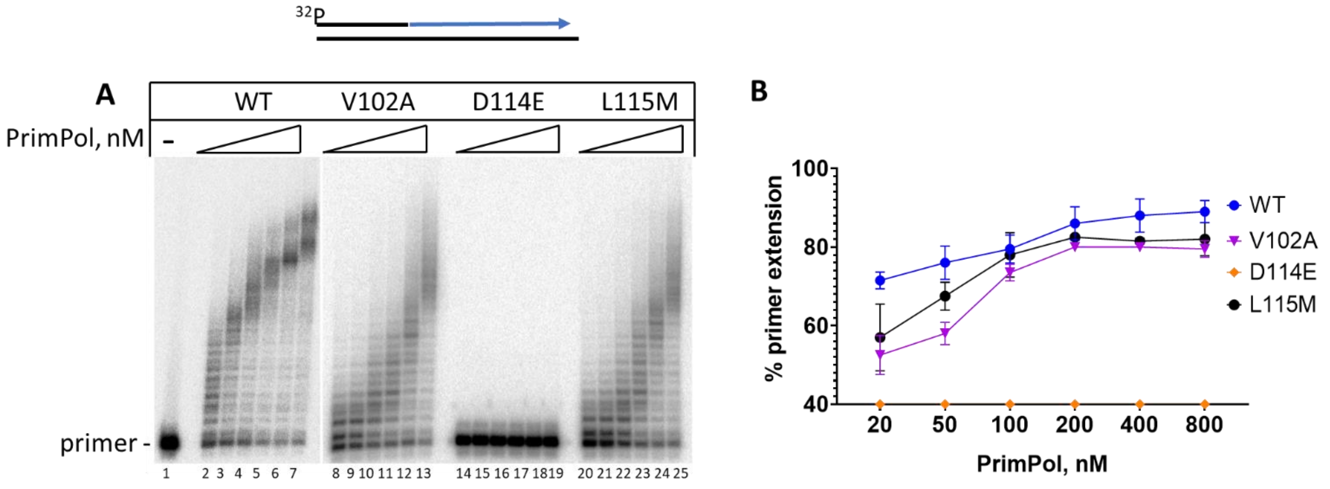


**Figure S2.** The DNA-polymerase activity of the wild-type PrimPol and its variants in the presence of $Mn^{2+}$ ions. **A**. The gel image showing primer extension by the wild-type PrimPol and its V102A, D114E and L115M variants on undamaged primer-template DNA substrate. Reactions were carried out in the presence of 1 mM $MnCl_2$, 200 μM dNTPs, 20/50/100/200/400/800 nM PrimPol, 20 nM $^{32}P$-labeled substrate for 10 min **B**. The graphs displaying dependence of DNA synthesis on protein concentration. Experiments were repeated three times, the mean values and standard errors are presented.

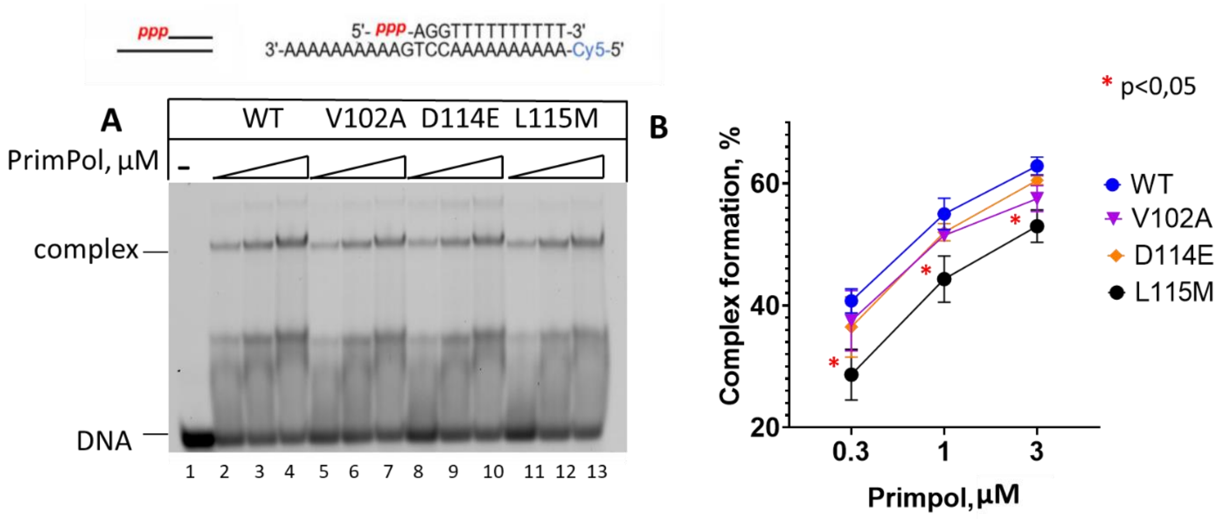


**Figure S3**. EMSA of binding to DNA substrate with the 5′-triphosphate by the wild-type PrimPol and its variants. **A**. Gel image showing the PrimPol-DNA complex formation. Reactions were incubated with 0.3/1/3 µM PrimPol in the presence of 300 nM DNA and 1 mM $Mn^{2+}$, pH 7.0. **B**. The graph showing the efficiency of the PrimPol-DNA complex formation by the wild-type PrimPol and catalytic domain variants. The experiments were repeated 2 – 4 times, the mean values and standard errors are presented.

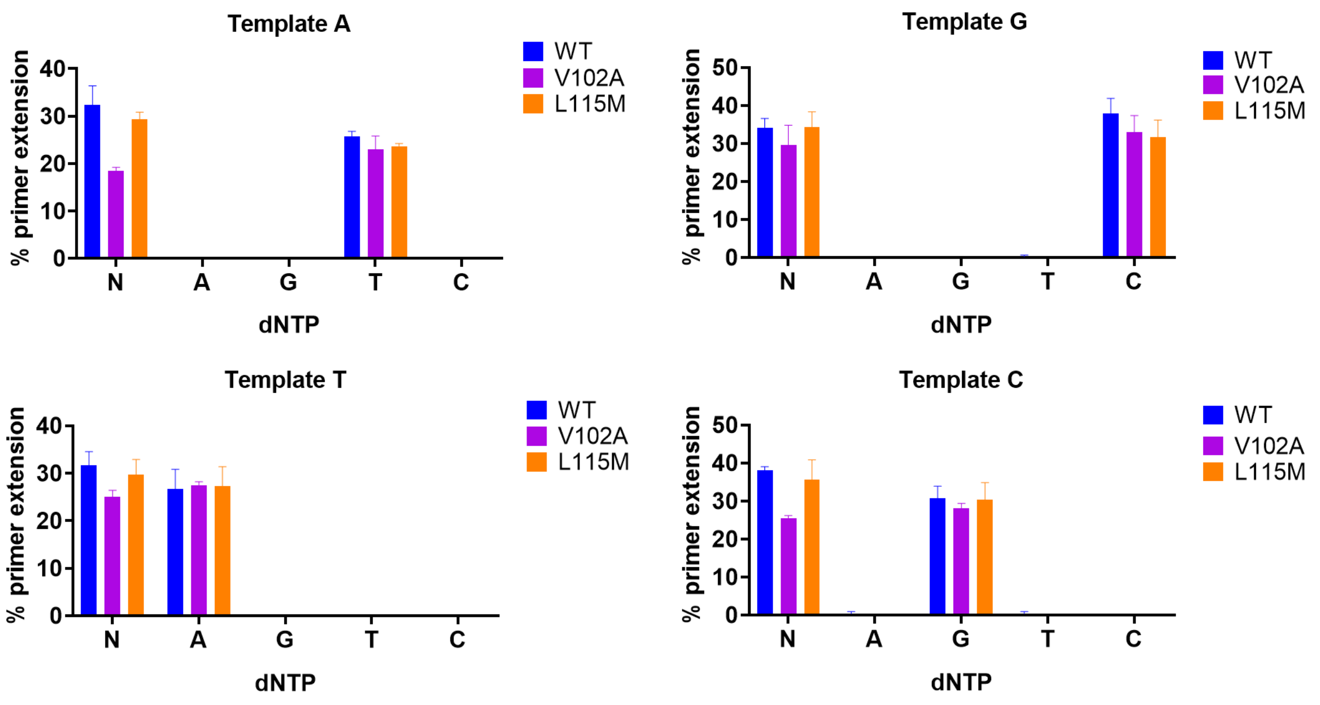


**Figure S4.** Individual dNMPs incorporation by PrimPol variants in the presence of $Mg^{2+}$ ions. The diagram showing the percentages of single dNMP incorporation of the wild-type PrimPol and its variants in the presence of $Mg^{2+}$ ions. The reactions were carried out in the presence of 1 mM $MnCl_2$, 100 nM Cy5-DNA substrate, 200 µM of dNTP or individual dATP, dGTP, dTTP, dCTP and 200 nM wild-type PrimPol (5 min), 600 nM V102A variant (90 min), 200 nM L115M variant (45 min). The experiments were repeated 2 – 4 times, the mean values and standard errors are presented.

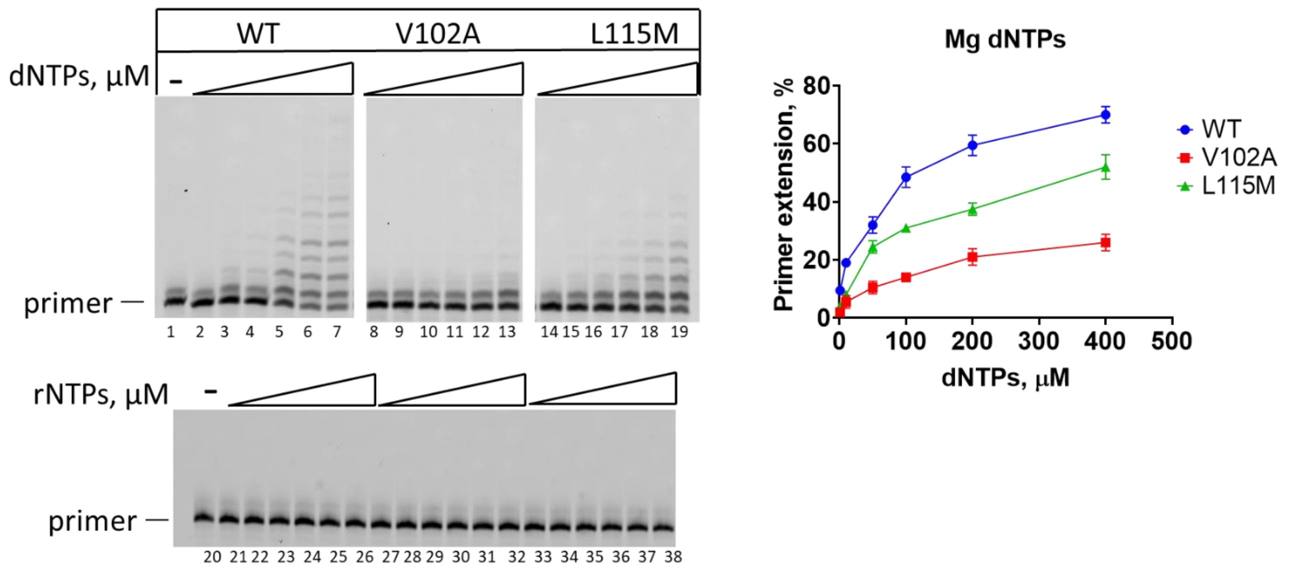


**Figure S5.** Ribonucleotides incorporation by PrimPol and its V102A and L115M variants in the presence of $Mg^{2+}$ ions. Reactions were carried out with 100 nM Cy5-labeled primer-template DNA substrate, 1/10/50/100/200/400/800 μM dNTPs or rNTPs, 200 nM wild-type PrimPol (20 min), 600 nM V102A variant (90 min), 400 nM L115M variant (40 min), 10 mm $MgCl_2$, pH 7.0.
